# CoReg, tracking-based motion compensation for *in vivo* calcium imaging

**DOI:** 10.64898/2026.11.09.763563

**Authors:** Luisa Neubig, Moritz Palm, Alessio Attardo, Joseph C. Donovan, Daniil Markov, Peter Soba, Neena Dihman, Shinjini Ghosh, Peter Rupprecht, Andreas M. Kist

## Abstract

In vivo calcium imaging enables the observation of neural activity with high spatial and temporal resolution. However, recordings are frequently affected by motion artifacts caused by animal movement, physiological processes, tissue drift, or instability of the imaging setup. These artifacts make signal extraction harder and reduce the consistency of downstream analyses. Current registration methods can correct rigid and non-rigid motion, but often estimate motion from individual frames or image patches and make limited use of temporal information. In addition, activity-driven fluorescence changes can make it difficult to distinguish changes in image appearance from actual motion.

Here, we treat motion correction in calcium imaging as a point tracking problem and introduce **CoReg**, an open-source Python-based software tool with a graphical user interface. We benchmark **CoReg** against established registration methods using fluorescence imaging datasets from multiple species. **CoReg** performs competitively with or better than existing approaches across several settings. We further show that its performance is robust to the choice of tracked points, making the method straightforward to apply. **CoReg** includes implementations of both our method and the evaluated established methods. Our results show that tracking-based registration provides a simple and reliable approach for motion correction in vivo calcium imaging.

## Introduction

In vivo calcium imaging is a widely used technique to study neural activity across different animal species and provides high spatial and temporal resolution^1–3^. The animal is generally spatially fixed to minimize movement during imaging. However, the degree of constraint and rigidity is often limited, for instance by the desire to simultaneously record animal behavior^4^. Hence, motion artifacts remain common and represent a major challenge for downstream processing. They can be caused by animal movement, physiological processes, slow tissue drift, or instability of the imaging setup. These artifacts impair signal extraction, reduce reproducibility, and complicate comparisons across experiments. Motion artifacts can occur in different forms. Linear motion mainly corresponds to translations of the field of view and can often be corrected using established registration methods. However, most microscopes acquire images by scanning, meaning each line or pixel is recorded at slightly different points in time. As a result, continuous linear motion during acquisition can produce non-linear distortions within a single frame, as different parts of the image reflect different positions of the sample^5^. Non-linear motion is more challenging because different image regions can move independently relative to each other and may undergo combinations of translation, shear, scaling, or local compression. These deformations, therefore, require more flexible correction methods. Motion within the imaged plane can, in principle, be corrected by any sufficiently flexible registration method. Motion out of the imaged plane, however, such as displacement along the Z-axis during an XY scan, removes tissue from the field of view entirely and cannot be recovered regardless of the correction method applied^6^. Current registration approaches typically correct each frame by computing its displacement relative to a fixed reference image or an iteratively updated template. While these methods can correct rigid and non-rigid motion, they often make only limited use of temporal information. This can be problematic in longitudinal calcium imaging, where motion develops continuously over many consecutive frames. In addition, fluorescence intensity changes are due to neural activity and not only due to motion, which can make intensity-based registration more difficult. Here, we propose to treat motion correction as a point tracking problem. Rather than estimating a deformation field for each frame independently, we select a sparse set of image points in the reference frame and track their positions across the full sequence. At each time point, the tracker infers positions by jointly processing a window of surrounding frames, rather than comparing individual frames in isolation. This produces trajectories that are smooth and consistent over time. We hypothesize that using this temporal information improves the robustness of motion estimation and enables the correction of both linear and non-linear motion artifacts in calcium imaging data. To test this, we introduce **CoReg** and benchmark it against established methods across fluorescence imaging datasets from multiple species. We further investigate how different point selection strategies affect registration performance. **CoReg** is released as an open-source tool that includes both our method and the evaluated established methods, allowing users to directly compare and explore different motion correction approaches.

### Related Works

Motion artifacts are a known problem in fluorescence microscopy, and commonly used methods register individual frames or image patches to a reference image or an updated template. StackReg^7^ in Fiji^8^, NoRMCorre^6^, Suite2p^9^, CaImAn^10^, and SIMA^11^ follow this general idea. NoRMCorre, for example, divides the image into overlapping patches and estimates local translations. Suite2p also uses block-wise registration to correct for non-rigid motion. Other methods, such as Moco^12^, focus on fast rigid registration. These frame-to-template approaches build on rigid registration methods originally developed for neuroimaging^13,14^. These methods work well in many settings, but motion is primarily estimated from frame-to-frame or frame-to-template comparisons. Temporal information is therefore used only to a limited extent. In calcium imaging, motion can develop gradually over many frames. If every frame or patch is corrected separately, the resulting deformation can change between consecutive frames and become temporally inconsistent.

Another challenge is that image intensities in calcium imaging are not constant over time. Fluorescence at any given pixel can change because of neural activity and not only because of motion. This makes it difficult to separate true motion from activity-driven intensity changes. Intensity-based registration methods can therefore estimate displacement from changes that are not caused by motion. Optical flow methods face a similar problem. They estimate dense displacement fields from intensity changes between frames^15,16^. Methods such as Flow-Registration^17^ have been developed for non-rigid motion correction in two-photon imaging. PatchWarp^18^ also models local deformations and can represent translation, rotation, scaling, and shearing. These approaches allow for more flexible corrections, but they still depend on appearance changes between images. More general non-rigid registration methods from medical imaging, such as LDDMM^19^ and ANTs^20^, can model complex deformations^21^. Learning-based approaches such as VoxelMorph^22^ can estimate deformations directly from data, but require training examples that may not be available for specialized microscopy modalities. However, they were mainly developed for anatomical images, where image intensities are more stable over time. In functional fluorescence imaging, changing activity patterns can make the estimation of deformation more difficult. Feature-based methods provide another way to estimate motion. Instead of using all image intensities, they rely on specific image structures or keypoints. Classical features such as SIFT, ORB, and AKAZE have already been investigated for two-photon calcium imaging^23^. These methods can be less sensitive to global intensity changes, but their performance depends on whether stable features can be detected in the image.

Tracking-based methods go one step further by following image structures over time. Aghayee et al.^24^ tracked neuronal particles across frames and used their trajectories for motion correction in 2D and 3D two-photon imaging. Landmark tracking has also been used to estimate tissue motion in other fluorescence imaging applications^25,26^. These studies show that sparse trajectories can provide useful information for motion correction. However, previous tracking-based approaches mainly rely on application-specific features or conventional tracking methods. Recent point tracking models such as CoTracker^27,28^ can track arbitrary points over long video sequences and jointly model several trajectories. This makes them interesting for motion correction because they track the same physical locations consistently across all frames, whereas frame-by-frame methods estimate correspondences independently and can produce inconsistent solutions over time. Existing methods have also rarely been evaluated across the full diversity of fluorescence imaging data. Recordings differ in species, preparation, motion type, fluorescence dynamics, signal-to-noise ratio, and acquisition settings. Most methods are evaluated on a limited number of datasets or specific experimental conditions. This makes it difficult to know how well a method transfers to other imaging scenarios.

Existing methods treat motion correction either as a frame-to-template alignment problem or as a dense optical flow problem. Neither approach uses temporal context across the full sequence. Neither exposes the estimated motion in an interpretable form. **CoReg** addresses both limitations by tracking a sparse set of points over time and recovering a dense deformation field from their trajectories. The resulting motion estimate is visible and editable by the user. We evaluate **CoReg** together with established methods across multiple fluorescence imaging datasets to investigate when tracking-based motion correction is reliable and useful.

In summary, we provide with **CoReg** the following contributions:

- Frame-wise registration metrics are insufficient surrogates for registration quality and should be complemented by downstream measures such as cell area preservation and Δ*F/F*_0_ fidelity
- A cell-based evaluation framework reveals that the choice of registration method affects cell morphology and signal fidelity
- **CoReg** allows users to inspect and adjust tracked points directly, making the correction process transparent and verifiable
- We release an open-source toolbox integrating **CoReg** and all baseline methods evaluated in this paper

## Materials and Methods

### Datasets

We evaluated **CoReg** on four calcium imaging datasets recorded from different species at different institutes using various microscopy techniques. The datasets comprise Danionella cerebrum, Drosophila melanogaster larvae, and two datasets from mouse (Mus musculus), one recording cortical activity using a wide-field head-mounted microscope and one recording dendritic activity using two-photon microscopy.

Calcium imaging data from Danionella cerebrum were acquired from a 5-month-old adult male expressing pan-neuronal, nuclear-localized GCaMP6s^29^, using oblique plane microscopy (OPM)^29,30^. All experimental procedures and animal husbandry complied with Berlin state, German federal, and European Union animal welfare regulations. The fish was immobilized as previously described^29^. It was anesthetized in 120 mg/L MS-222 and transferred to a custom agar mold. Oxygenated water was delivered through a pipette inserted into the mouth using a peristaltic pump. The fish was fixed in place with a few droplets of low-melting-point agarose (Fisher Scientific) and positioned under the OPM objective. Volumetric imaging was performed at 1 Hz for 20 minutes. Following volumetric reconstruction and de-shearing, three individual imaging planes were extracted without any motion correction or registration. A 10-minute segment was then extracted from these planes. This segment contained several motion artifacts induced by tail movements and characteristic periodic wobbling of the OPM images, potentially associated with gill movements.

Mouse data from dorsal hippocampal CA1 were acquired using a Wide-Field Head Mounted (WFHM) microscope^31^. Animals were C57BL/6 male mice between three and six months of age. The objective lens (0.5 NA, 2.4 mm WD, air immersion) and the microscope (Doriclenses, S model) were mounted on the head of each animal. Animals were free to explore an open arena for 5–10 minutes per day. Ca^2+^ transients were recorded for 15 minutes at a sampling rate of 45 Hz using a 488 nm continuous wavelength laser (Thorlabs) at 1 mW average power and Doric Neuroscience Studio software. All animal procedures conformed to the guidelines of the Max Planck Society and were approved by the local animal authority (Regierung von Oberbayern—Veterinärwesen) under the IRB #ROB-55.2Vet-2532.Vet_02-16-48.

Dendritic data were acquired from CA2/CA3 of mouse hippocampus using a two-photon resonant scanning microscope with a K-mirror for field-of-view rotation and an electrically tunable lens for curving the field of view along the slow scan axis, as described in^32^. Calcium imaging was performed at 1024 *×* 1024 pixels and a frame rate of 15 Hz. GCaMP8m expression was induced in a wild-type C57BL/6 mouse by stereotaxic injection of ssAAV-9/2-mCaMKII*α*-jGCaMP8m-WPRE-bGHp(A) (Viral Vector Facility Zürich) into hippocampus (AP *−*2.0 mm, ML *−*1.8 mm from bregma, DV *−*1.3 mm from brain surface). Two weeks after injection, a hippocampal window was implanted as previously described^33^. Two weeks after surgery, the animal was habituated to head fixation under the microscope. Ca^2+^ imaging was performed during spontaneous activity, with the animal seated in a 3D-printed chair in a perched posture^34^. All animal procedures were conducted in accordance with Swiss federal guidelines and approved by the Cantonal Veterinary Office of Zürich.

Drosophila larvae data were acquired to monitor neuronal responses to noxious heat stimulation. Calcium imaging was performed in intact third-instar Drosophila melanogaster larvae expressing GCaMP6s under a motor neuron-specific GAL4 driver (Hb9-GAL4). Fluorescence was recorded from neuronal somata within the larval ventral nerve cord. Individual third-instar larvae (92 *±* 4 h after egg laying) were mounted in a custom-built chamber with a local heat stimulator (infra-motion, Hamburg). Imaging was performed using a Zeiss LSM 900 AS2 confocal microscope equipped with a 20 *×*/NA 0.8 air objective. Noxious heat stimulation at 46°C was delivered to abdominal segments A3–A6 while calcium responses were recorded from the ventral nerve cord. Time-lapse recordings were acquired at 4.3 fps with a resolution of 512 *×* 258 pixels, collecting 750 consecutive frames per larva. Consistent acquisition settings were maintained across all recordings.

The four datasets cover a range of motion types. The Drosophila larvae dataset exhibited predominantly global non-linear motion, ranging from low to strong movement. The remaining three datasets showed more local or temporally shorter motion patterns. Together, they allowed us to assess the robustness of **CoReg** across different biological preparations and motion types.

### Quantitative and Qualitative Evaluation

The goal of motion correction is to align all frames in a calcium imaging recording to a common reference so that the apparent position of imaged structures remains consistent over time and can be analyzed in downstream tasks. We evaluate registration quality using five frame-wise metrics and three self-consistency metrics. All metrics are computed after percentile normalization of the video, using the 1st and 99th percentiles to scale intensity values to the range [0, 1]. Frames are also cropped by a border margin of 10% to make different border handling (e.g., zero filling or reference frame filling) comparable.

#### Mean squared error (MSE)

The mean squared error between the reference frame *r* and the current frame *c* is computed pixelwise and averaged:

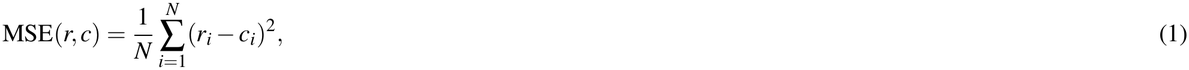

where *N* is the number of pixels and *r*_*i*_, *c*_*i*_ are the intensity values at pixel *i*. MSE measures overall pixel-level intensity similarity between frames. Lower values indicate better alignment. MSE is averaged across all frames.

#### Peak signal-to-noise ratio (PSNR)

The peak signal-to-noise ratio is derived from the MSE. Since intensity values are normalized to [0, 1], the maximum possible intensity is 1, and PSNR simplifies to:

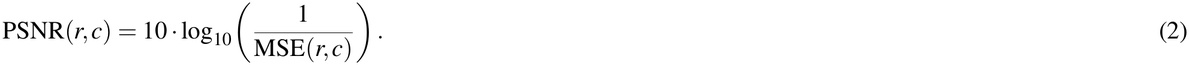

PSNR provides a logarithmic representation of MSE. Higher values indicate better alignment. PSNR is averaged across all frames.

#### Zero-normalized cross-correlation (ZNCC)

The zero-normalized cross-correlation is computed for each frame pair consisting of the reference frame *r* and the current frame *c*, and then it is averaged across all frames:

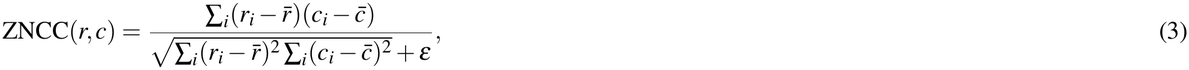

where 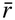 and 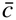 are the mean intensity values of the reference and current frames, respectively. This is equivalent to the Pearson correlation coefficient between pixel intensities. A value of *ε* = 10^*−*12^ is added to avoid division by zero. ZNCC measures the linear intensity correlation between frames, invariant to global differences in mean and scale. The self-comparison of frame 0 is fixed to 1.0.

#### Structural similarity index measure (SSIM)

The structural similarity index measure^35^ captures luminance, contrast, and structural differences between the reference frame *r* and the current frame *c*:

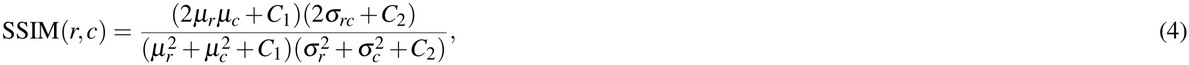

where *µ*_*r*_, *µ*_*c*_ are the local means, 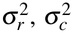 the local variances, and *σ*_*rc*_ the local covariance of *r* and *c. C*_1_ and *C*_2_ are small constants added for numerical stability. SSIM evaluates the preservation of local image structure by comparing luminance, contrast, and spatial intensity patterns between frames. SSIM values range from *−*1 to 1, with 1 indicating perfect agreement. SSIM is averaged across all frames.

#### Mutual information (MI)

Mutual information measures the statistical dependency between the intensity distributions of the reference frame *r* and the current frame *c*:

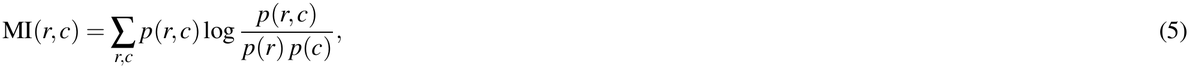

where *p*(*r, c*) is the joint intensity histogram and *p*(*r*), *p*(*c*) are the marginal histograms, estimated from the respective frame. MI captures image similarity even when the intensity relationship between frames is nonlinear. Higher values indicate greater statistical dependency and better alignment. MI is averaged across all frames.

#### Crispness (Crisp)

Crispness is computed from the temporal mean image *Ī* and not separately for each frame:

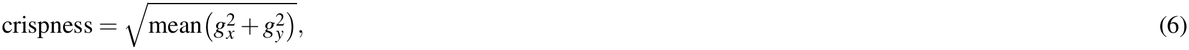

where *g*_*x*_ and *g*_*y*_ are the spatial gradients of the temporal mean image in the horizontal and vertical directions, computed using `numpy.gradient`. Crispness reflects how well sharp image structures are preserved across the recording. Higher values indicate a sharper mean image and, therefore, less motion blur caused by misalignment. Poor registration results in a softer mean image with a lower gradient magnitude.

#### Correlation with the temporal mean image (CorrMean)

The mean correlation corr_mean_ is calculated using the same Pearson correlation formulation as ZNCC. Each frame is correlated with the temporal mean image *Ī* rather than with frame 0, and the values are averaged across all *T* frames:

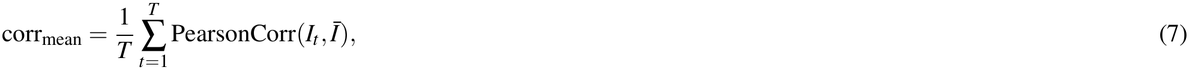

where *I*_*t*_ denotes the frame at the time point *t*. CorrMean reflects the temporal consistency of registered frames relative to the overall mean appearance of the video. Higher values indicate better alignment.

#### Residual optical flow (ResFlow)]

The temporal mean image is used as a reference template. TV-L1 optical flow^36^, implemented via `skimage.registration.optical_flow_tvl1`^37^, is computed between the temporal mean image and each frame. For each sampled frame, the residual flow is the median magnitude of the dense displacement field:

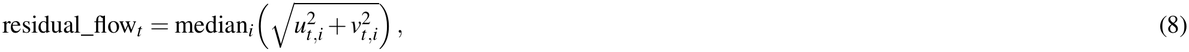

where *u*_*t,i*_ and *v*_*t,i*_ are the horizontal and vertical components of the optical flow at pixel *i* between the mean image and frame *t*. The final value is the mean of these across all sampled frames:

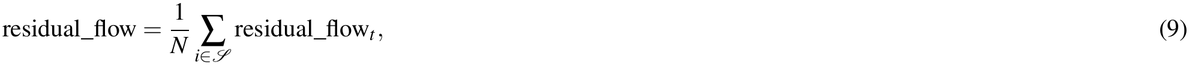

where *S* is the set of sampled frames and *N* = |*S*| . ResFlow directly quantifies how much spatial misalignment remains in the corrected video, independently of intensity changes caused by neural activity.

#### Downstream analysis

For the downstream evaluation, we annotated cells in the raw and motion-corrected data. To assess the consistency of extracted calcium signals, we measured the Δ*F/F*_0_ of activated neurons and the area of manually annotated cells. We compared cell activity in the raw video against the tracked raw video and the motion-corrected data. We also evaluated cell sizes between the motion-corrected and raw data to assess how each method affects cell size.

### CoReg

CoReg is a versatile motion correction tool for fluorescence microscopy. Unlike algorithmic, black-box-like methods such as ANTs, NoRMCorre, or PyFlowReg, CoReg uses a set of tracking points that users can inspect and modify. They can add, remove, or manually correct these points before and after registration. This makes the correction process easier to understand and verify. Users can identify which regions contribute to the estimated motion, reduce the number of points to simplify the deformation model, and correct inaccurate tracks in difficult cases. We treat motion correction as a point tracking problem. Rather than registering full frames to a reference, we select a set of points in the reference frame and track their positions across all subsequent frames by using CoTracker3^28^ as the underlying tracking algorithm in an offline fashion, meaning tracking is verified both forwards and backwards in time. CoTracker3 is a transformer-based model that estimates the positions of multiple points across a sliding temporal window of frames. Instead of tracking each point in isolation, it uses attention across all tracked points simultaneously, allowing nearby points to share motion information. This co-tracking approach improves robustness in low-contrast regions and under fluorescence changes that can resemble motion. The key advantage over dense patch-based registration is temporal consistency. Because all points are tracked across the full sequence in one pass, the resulting trajectories are temporally consistent by construction. Frame-by-frame methods, by contrast, estimate each deformation independently and can produce inconsistent solutions between frames. Points are selected sparsely in regions where tracking is reliable, and a smooth, dense deformation field is recovered from these sparse trajectories via thin-plate spline interpolation^38^, which is sufficient because biological tissue motion is typically spatially smooth. Regions outside the field of view after transformation are filled with zeros. On a single GPU (RTX A5000), processing 500 frames with a single preset, including tracking and warping, takes approximately 3–4 minutes. An overview of the framework is shown in Figure 1. CoReg can be used through a command-line interface or an interactive graphical user interface. An overview of the graphical user interface is shown in Supplementary Figure 1 and how it is used is documented in Supplementary Figure 2 and Supplementary Movie 1. Tracks can be saved and exported to allow interoperability and integration into existing workflows.

**Figure 1.**
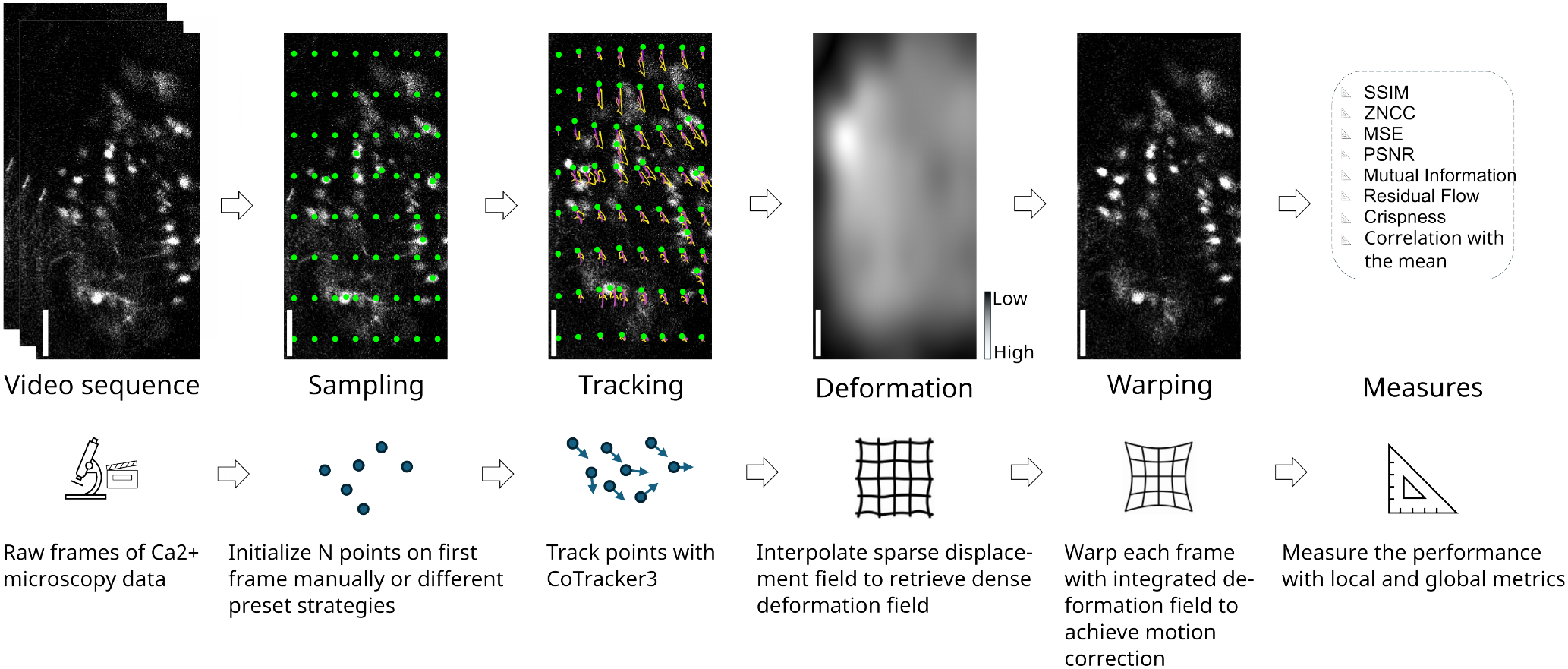
**CoReg** framework overview. Given a video sequence, sampled points are tracked across time using the CoTracker framework. Deformation fields are computed through the tracked points and used to warp the respected frames. In this study, we assessed the registration performance across eight different local and global measures. Scale bar indicates 50 µm.

CoReg provides six automatic methods for selecting tracking points. The available presets and their combined usage are shown in Supplementary Figure 3:

- **Regular grid:** Points are placed on a uniform grid spanning the full frame. The grid is defined by a row and column count and can be shifted by a pixel offset. A fixed-spacing variant constructs the same grid from a target pixel spacing instead.
- **Features:** Shi-Tomasi corners are detected using `goodFeaturesToTrack` from OpenCV. Detection is restricted to the foreground, defined by Otsu thresholding. This favors high-contrast structural landmarks.
- **Motion:** Points are placed where the unregistered image changes most between frames. A motion map is built from the summed absolute difference between frames and then thresholded into connected components. Each component that exceeds a minimum motion strength receives at least one point. Remaining points are distributed in proportion to each component’s motion.
- **Greedy motion:** This strategy also uses the inter-frame motion map but distributes points more evenly. The motion map is smoothed and compressed into a sampling density. Seed points are placed at local peaks of this density, and additional points are drawn in proportion to it while maintaining a minimum distance between them. This provides better spatial coverage than *Motion* while still focusing on active regions.
- **Segmentation:** The frame is thresholded using Otsu’s method into connected components. One point is placed at the centroid of each of the largest components.
- **MatchAnything:** Points are obtained from a pretrained keypoint-matching model^39^. The reference frame is matched against a second frame, and only the highest-confidence, well-separated matches are kept. If the model is not available, the method falls back to the *Features* strategy.

### Baseline Methods

We compare **CoReg** against NoRMCorre^6^, ANTs^20^, LDDMM^19^, and PyFlowReg^17^. All four are also integrated into the **CoReg** application. All methods ran on a GPU for evaluation (PyTorch 2.5.1, CUDA 12.1).

NoRMCorre was run through jnormcorre (v1.0.0), a JAX-accelerated re-implementation of the original method^40^. We used the piecewise-rigid variant. All other parameters match the jnormcorre defaults: maximum shifts of 6 pixels, strides of 96, overlaps of 32, FFT upsampling factor of 4, and maximum rigid deviation of 3.

ANTs was run through antspyx (v0.6.3)^41^, the official Python bindings for the ANTs C++ library. We used SyNOnly instead of the default SyN transform, which omits the initial rigid and affine alignment steps to focus the evaluation on non-rigid deformation correction.

PyFlowReg was run through the PyFlowReg package (v0.1.0a9). Alpha was set to 1.5, matching the package default. The minimum pyramid level was set to 4.

LDDMM was run through an in-house PyTorch implementation using velocity-field parameterization, a Gaussian-kernel smoother, and a gradient-descent optimizer with a reducing step size because no maintained pip-installable LDDMM package exists. Parameters were tuned manually: kernel width *a* = 40, *n*_*t*_ = 5 time steps, 100 gradient-descent iterations, step size *ε* = 0.001, matching-term weight *σ* = 0.1, regularization weight *σ*_*R*_ = 1.0. An initial affine alignment step preceded the diffeomorphic step.

## Software and Data availability

CoReg is available as an open-source software package under the MIT License at https://github.com/ankilab/CoReg. The package includes the proposed method and all baseline methods evaluated in this study (NoRMCorre, PyFlowReg, ANTs, and LDDMM). The software is implemented in Python 3.12.13. The graphical user interface is built with PyQt6. Motion compensation is performed using PyTorch 2.5.1 with CUDA 12.1. Point tracking is based on the CoTracker offline model (v3). An additional Napari plugin makes motion compensation with all presets available directly within the image viewer.

## Results

### Motion artifacts and correction examples

Motion artifacts in calcium imaging vary in their nature, and it remains unclear whether a single method can reliably correct both linear and non-linear deformations. In contrast to existing methods, we investigated the strength of tracking-based motion estimation of pixels and their underlying structures. Although **CoReg** uses a tracking model that is not trained on biomedical data, it already produces high-quality tracking results for very noisy calcium imaging data (see Supplementary Movie 2). The deformation field crucial for registration is generated through the individual point tracks set in the video sequence (Fig. 1 and Methods). We reasoned that point placement is a key determinant for tracking and the resulting motion compensation. Therefore, we evaluated a variety of different point selection strategies that are also implemented as presets in the **CoReg** toolbox. In total, we evaluated six methods, namely Features, Regular grid, Segmentation, Motion, MatchAnything, and Greedy motion. Some are data agnostic (Regular grid), some are motion-driven (Motion, Greedy motion), and some are structure-driven (Features, MatchAnything, Segmentation). For our experiments, we used a 16 *×* 16 grid for the Regular grid and an upper limit of 500 points for the other structure-driven point presets. We chose the 16 *×* 16 grid based on a systematic evaluation of grid density. Beyond a certain number of points, adding more grid points does not improve the quality of the recovered deformation field. The 16 *×* 16 configuration was identified as the point where performance plateaus, making it the most efficient choice for our experiments (Supplementary Figure 4, Supplementary Movie 3). A systematic evaluation of grid density is also provided in Neubig et al.^42^.

For our first analysis, we used the Drosophila larvae dataset, as it contains recordings with both strong and low movement, allowing us to assess performance across different motion conditions. In Figure 2a, we show the PSNR, ZNCC, CorrMean, and ResFlow for the **CoReg** pipeline with six different point sampling approaches. These four metrics are representative of the overall trends observed across all eight metrics. Results for the remaining metrics are provided in Supplementary Figure 5. For all four metrics, the results seem almost equal for the low movement cases, whereas Greedy motion shows worse results on the strong-movement data. For the later analysis, we only use **CoReg** with the *Regular grid* as it showed the most consistent results. Next, we were interested in whether tracking-based registration is comparable to established methods. Therefore, we compared **CoReg** (Regular grid) against the established methods: ANTs, NoRMCorre, PyFlowReg, LDDMM, and evaluated it for PSNR, ZNCC, CorrMean, and the ResFlow (Figure 2b). We found that **CoReg** consistently outperforms ANTs, NoRMCorre, and LDDMM across all metrics, regardless of low or strong movement. PyFlowReg, however, achieves results equal to or slightly better than **CoReg** in some of the analyzed metrics. A key advantage of **CoReg** is its ability to track corresponding points across long image sequences, producing temporally consistent motion estimates. We therefore tested whether this long-term tracking provides more reliable motion estimates than the frame-wise deformation information used by the baseline methods. Here, we compare our tracking-based method against PyFlowReg, which uses an optical-flow-based approach with a registration of each frame to a common reference image, whereas ANTs and LDDMM use numerical deformation mapping, and NoRMCorre performs piecewise-rigid template matching by aligning overlapping image patches to an iteratively updated reference template. We compared all five methods for sequences of 50, 100, 250 frames, and the full sequence (*∼* 760 frames), as shown in Figure 2c. ANTs, LDDMM, and NoRMCorre performed worst in SSIM and ZNCC for both cases. PyFlowReg shows slightly higher accuracy in motion compensation for SSIM and ZNCC. Overall, **CoReg** performs at least on par with PyFlowReg across all sequence lengths, while the other baseline methods show a stronger decline in performance as the sequence length grows. Frame-wise registration metrics capture spatial alignment quality but may not fully reflect whether motion has been genuinely corrected. Temporal standard deviation (std) provides a complementary measure. Motion increases pixel-level intensity fluctuations over time, so a method that effectively removes motion should lower the std relative to the unregistered video. Mean image sharpness offers a further check, as inaccurate deformation fields introduce blur. We therefore analyzed both measures for the Drosophila larvae dataset (Figure 3). At low movement, **CoReg** presets achieved the strongest std reduction overall. At strong movement, **CoReg** and PyFlowReg produced comparable std reductions, both substantially lower than the remaining established methods (Figure 3b). **CoReg** and PyFlowReg reached the lowest temporal std (0.18 *×* and 0.19*×* raw) and the highest sharpness (2.45 *×* and 2.30 *×* raw). LDDMM, by contrast, reduced the std only modestly (0.75 *×* raw) and produced blurred mean images with sharpness falling below the unregistered baseline (0.87*×* raw), indicating that its deformations displaced rather than corrected image content.

**Figure 2.**
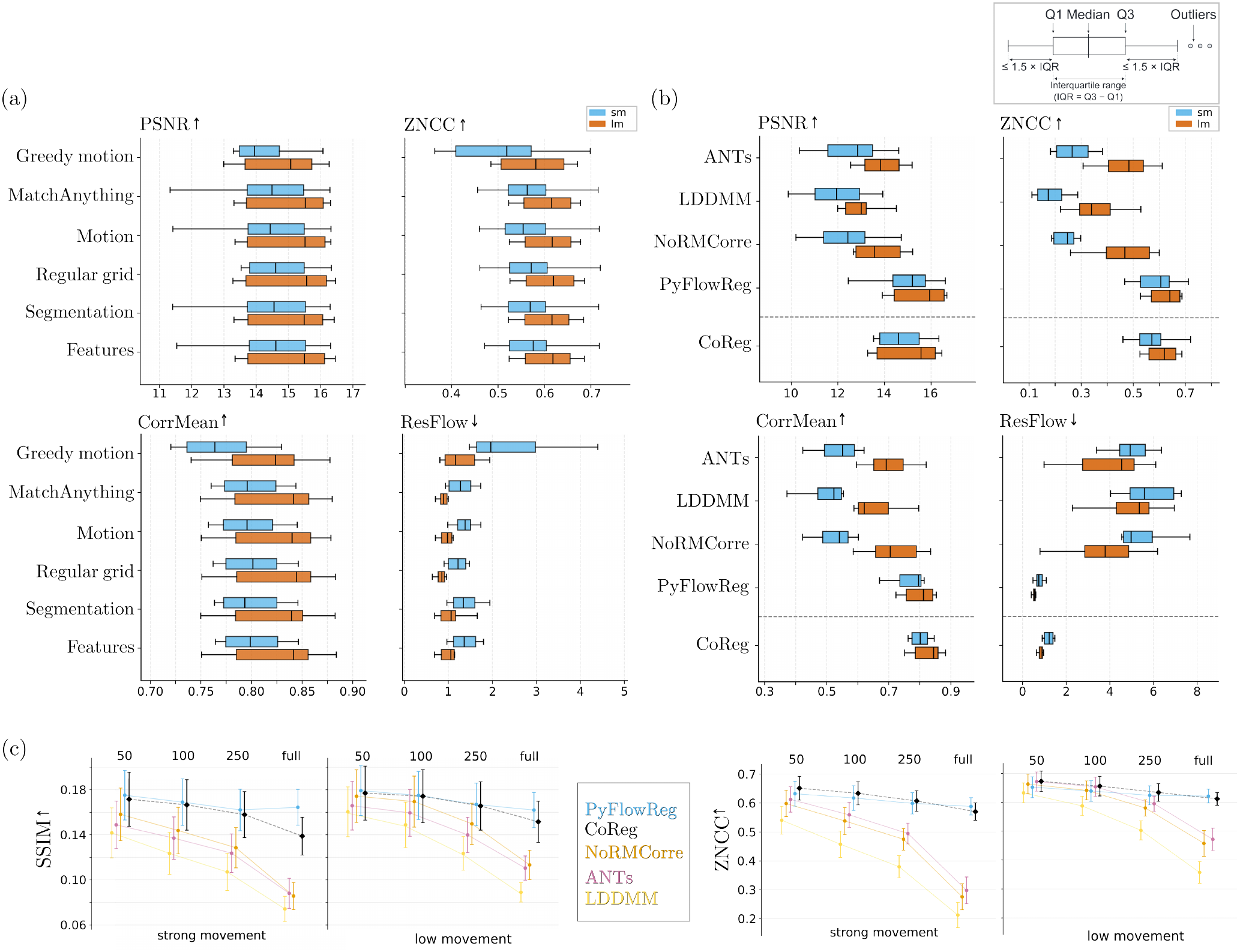
CoReg framework evaluation for the Drosophila larvae data. Box plots show the median (center line), interquartile range (box), and 1.5*×* IQR whiskers; individual outliers are shown as points. (a) Differences in 4 of 8 metrics of the initial point sets for the **CoReg** algorithm. Recordings were split into eight strong-movement (sm; blue) and eight low-movement (lm; orange) recordings. (b) Metric-wise comparison of the baseline methods (ANTs, LDDMM, NoRMCorre, PyFlowReg) against **CoReg** (Regular grid). (c) Influence of sequence length on motion compensation (established methods vs. **CoReg** (Regular grid)).

**Figure 3.**
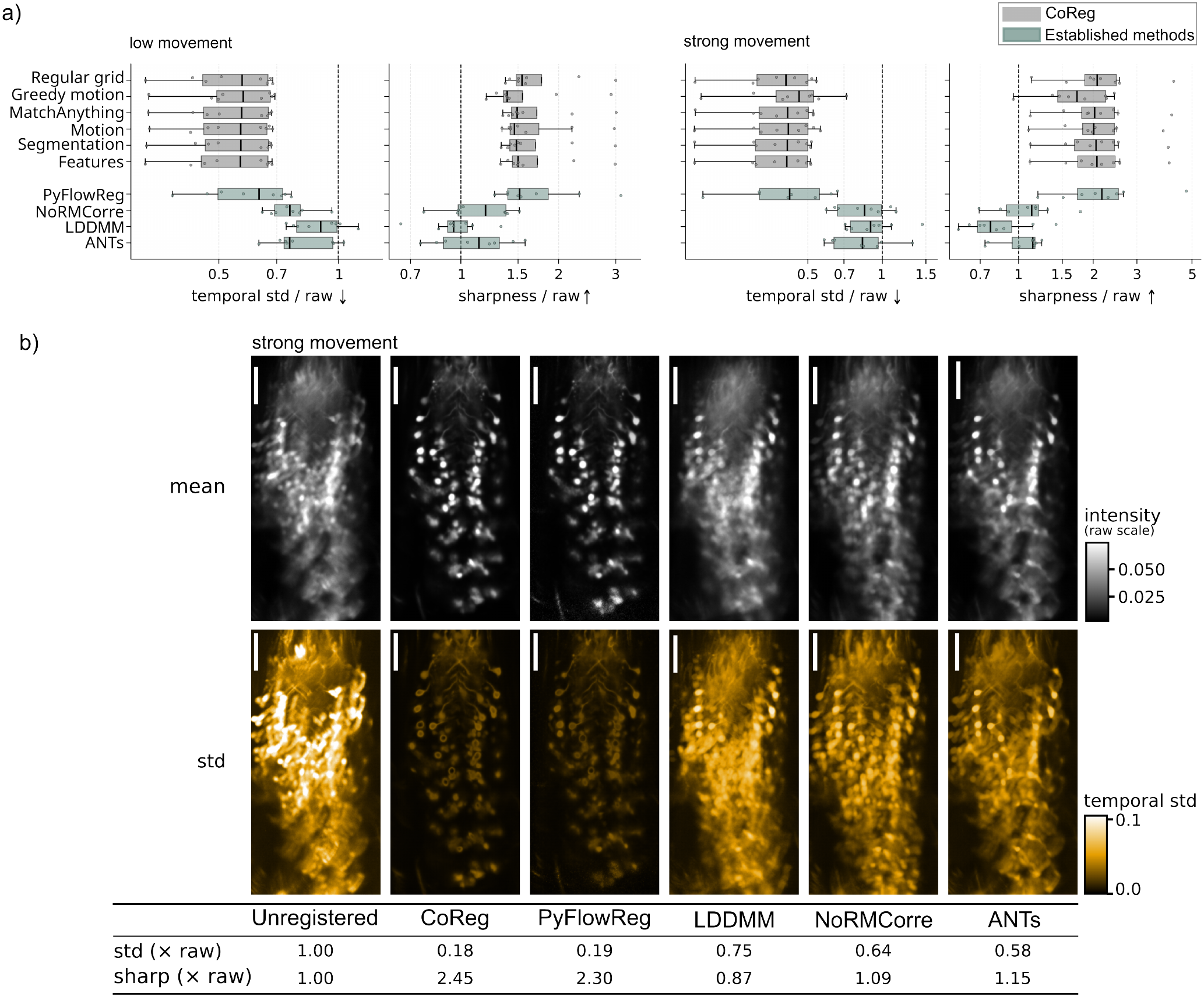
Temporal standard deviation and image sharpness analysis for the Drosophila larvae dataset. (a) Temporal standard deviation (std) and mean image sharpness relative to the unregistered video for all **CoReg** presets (light gray) and established methods (light green-gray), across low-movement (*N* = 8) and strong-movement (*N* = 8) recordings. A ratio below 1 indicates a reduction and a ratio above 1 indicates an increase relative to the unregistered video. The dashed line marks the unregistered baseline. (b) Representative strong-movement recording. The top row shows mean projection images and the bottom row shows temporal std images. The table lists std and sharpness values relative to the unregistered video for each method. Scale bar indicates 50 µm.

### Effects on downstream analysis

The goal of motion correction is to improve the quality of cellular signals extracted in downstream analyzes while minimizing distortions of the underlying data. Our quantitative evaluation showed that CoReg and PyFlowReg achieve comparable performance across standard registration metrics. However, these metrics primarily assess image similarity and may not fully capture how well the original signal information is preserved. We therefore complemented the quantitative comparison with a qualitative assessment of the registered images, focusing on how well each method compensated for motion while preserving the original image content. We analyzed overlays of the first and last frames for each method (Figure 4a). White areas indicate cells in the reference frame. In a well-aligned result, colored cells overlap with these white areas, with any residual color confined to their immediate surroundings. Colored areas in the background between cells suggest misalignment. CoReg and PyFlowReg showed good overall alignment, while NoRMCorre, LDDMM, and ANTs showed the least agreement between the reference and registered frames. To examine this in more detail, we cropped a representative region shown in Figure 4b. NoRMCorre, LDDMM, and ANTs show little overlap between white and colored cells (dark orange arrowheads), indicating that residual drift remains. CoReg shows the expected pattern, with colored regions concentrated around white cell outlines (green arrowheads). PyFlowReg also achieves good cell alignment (green arrowheads) but shows colored areas spreading into regions between cells (dark orange arrowheads), suggesting incomplete alignment in those regions. During qualitative inspection, we noticed that some methods visibly altered cell size in the corrected data. To quantify this, we annotated 3 cells in 6 single planes over 30 frames each (18 cells in total) and evaluated the ratio of the cell size in the corrected video to the size in the original video in the same 30 frames. A ratio of 1 indicates that the original cell size is preserved. We present the size ratio across all cells for all five methods in Figure 4c. **CoReg** shows the most consistent mean and preserves the original cell size most accurately. PyFlowReg, NoRMCorre, and LDDMM report a smaller cell size than the original, while ANTs tends to expand it. As **CoReg** already demonstrates the best preservation of cell size, we asked whether this is also reflected in the maintenance of cell activity over time. We evaluated five cells in 16 stacks and computed the Δ*F/F*_0_, estimating *F*_0_ as the mean of the first five frames of each cell in the unregistered video. Figure 4d shows one example where the ROIs are drawn into the video (see Supplementary Movie 4). NoRMCorre, LDDMM, and ANTs completely lose the cell due to poor motion compensation, resulting in near-zero signal amplitude. Both PyFlowReg and **CoReg** correctly outline the cell and recover calcium transients. However, **CoReg** most accurately preserves the signal amplitude relative to the unregistered tracked ROI, while PyFlowReg shows lower peak Δ*F/F*_0_ values, consistent with the cell size underestimation observed above.

**Figure 4.**
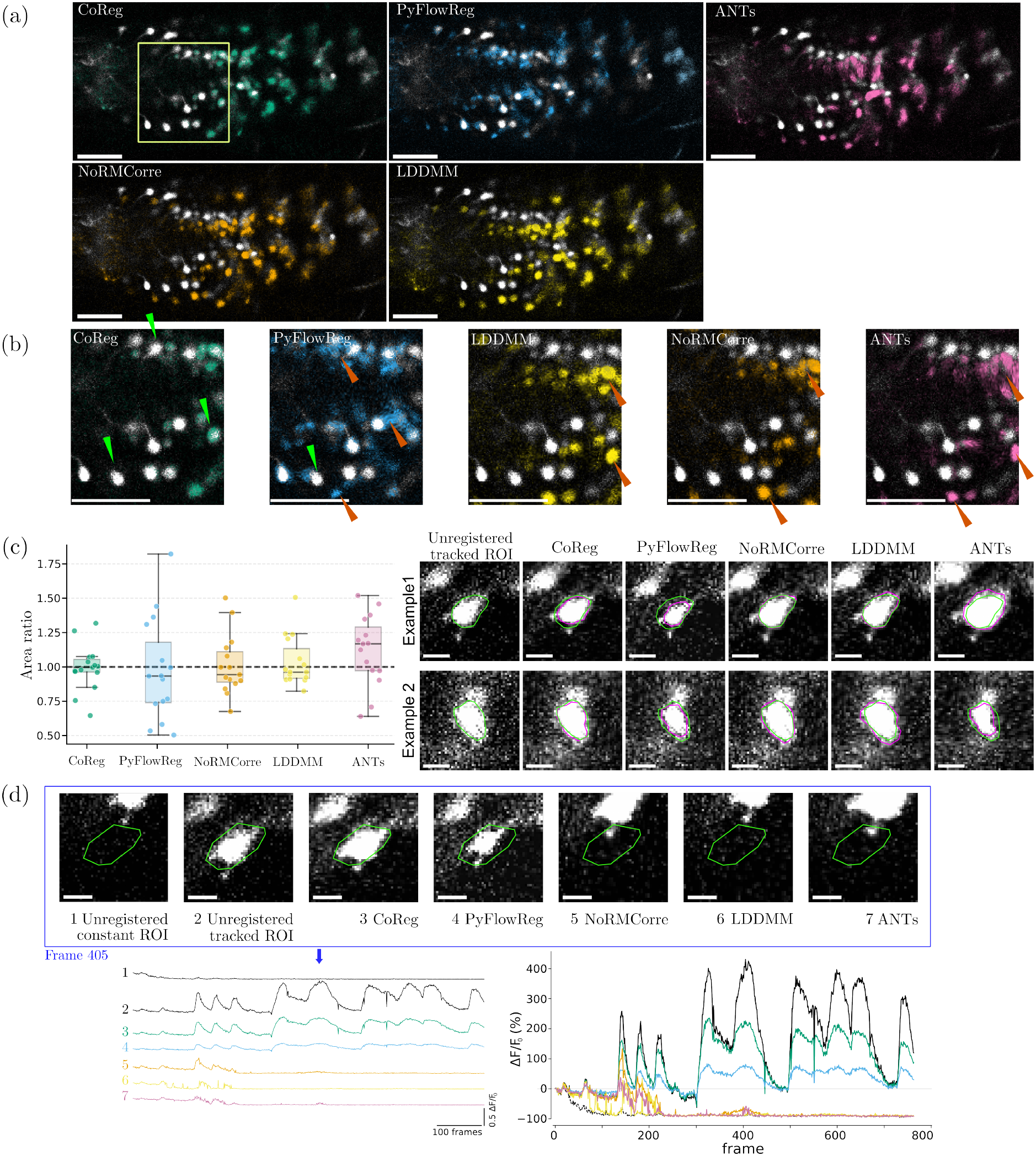
**CoReg** preserves cellular signal and cell morphology for downstream analysis. (a) Overlay of the first and last frame for each method. White areas indicate cells in the reference frame. In a well-aligned result, colored cells overlap with these white areas, with any residual color confined to their immediate surroundings. Colored areas in the background between cells suggest misalignment. (b) Representative crop of (a) for each method, highlighting regions of alignment and misalignment. Green arrowheads indicate well-aligned structures. Dark orange arrowheads indicate alignment artifacts. Scale bar indicates 50 µm. (c) Quantitative analysis of the cell area ratio of the motion-compensated frames relative to the unregistered video (N=18). A ratio of 1 indicates perfect size preservation (dashed line). Box plots follow the same convention as Figure 2. Two qualitative examples show the unregistered cell outline in green and the registered cell outline in magenta for each method. (d) Extracted Ca^2+^ activity for an example cell across all methods. The upper panel shows the ROI (green outline) overlaid on a representative frame (frame 405, blue arrow) for each condition. The left plot shows the full Δ*F/F*_0_ traces across all frames for all conditions. The right plot shows the same Δ*F/F*_0_ traces on a common axis, allowing direct comparison of signal amplitude across methods. Scale bar indicates 10 µm.

### Generalizability across datasets

In order to test **CoReg**’s performance across imaging modalities and on different organisms, we applied **CoReg** to whole-brain OPM recordings of Danionella cerebrum and two-photon recordings of cortical activity in mice. Motion characteristics varied strongly across datasets. The Drosophila larvae recordings showed both linear and non-linear motion artifacts across the full sequence, ranging from low to strong movement. The mouse recordings showed shimmering motion. The Danionella whole-brain recordings exhibited motion artifacts spanning fewer frames (motion period < 10 frames). Across both datasets, **CoReg** presets and established methods performed comparably across most registration metrics (Supplementary Figure 6). For the Mice dataset, the Greedy motion preset showed the highest residual flow, indicating inconsistent motion estimates in some recordings. For the Danionella dataset, LDDMM was a consistent outlier, performing the worst across multiple metrics. As discussed above, frame-wise metrics may not fully capture the quality of motion compensation, so we applied the same temporal std and sharpness analysis to both datasets (Fig. 5). For the Danionella dataset, most methods lowered the std (see Fig. 5a). Among the established methods, ANTs performed best, reducing the std in almost all slices, while PyFlowReg changed it only slightly. LDDMM showed severe issues in the registration, with the median std above that of the unregistered data in several slices. For the mice dataset, most methods lowered the std, with the exception of the Greedy motion point selection preset, which failed in about half of the chunks, with its std sometimes exceeding the raw data by an order of magnitude. ANTs performed best overall, followed by several **CoReg** presets. Figure 5b shows two example recordings. In the mouse example (top), **CoReg** gave the lowest temporal std and increased mean image sharpness, while LDDMM, despite also reducing the std, introduced bright band artifacts visible in the std image, suggesting the image content was displaced rather than corrected. In the Danionella example (bottom), PyFlowReg and LDDMM increased the temporal std relative to the unregistered video, whereas **CoReg** and ANTs reduced it, and NoRMCorre left it unchanged. PyFlowReg achieved the highest sharpness in both examples (mice: 1.47*×*, Danionella: 1.24 *×* raw), but in the Danionella recording, it also increased the temporal std to 1.07 *×* raw, suggesting that the sharpness gain does not reflect improved motion compensation. The temporal std also contains contributions from neural activity and shot noise, so a ratio close to 1 does not indicate that a method had no effect. Across all three datasets, the best-performing **CoReg** presets consistently reduced the temporal std, demonstrating robust motion compensation across different biological preparations and motion types. We additionally applied **CoReg** to a dendrite imaging dataset. Point tracking produced reliable trajectories (Supplementary Movie 5) and motion compensation was qualitatively comparable across **CoReg**, PyFlowReg, and ANTs (Supplementary Movie 6).

**Figure 5.**
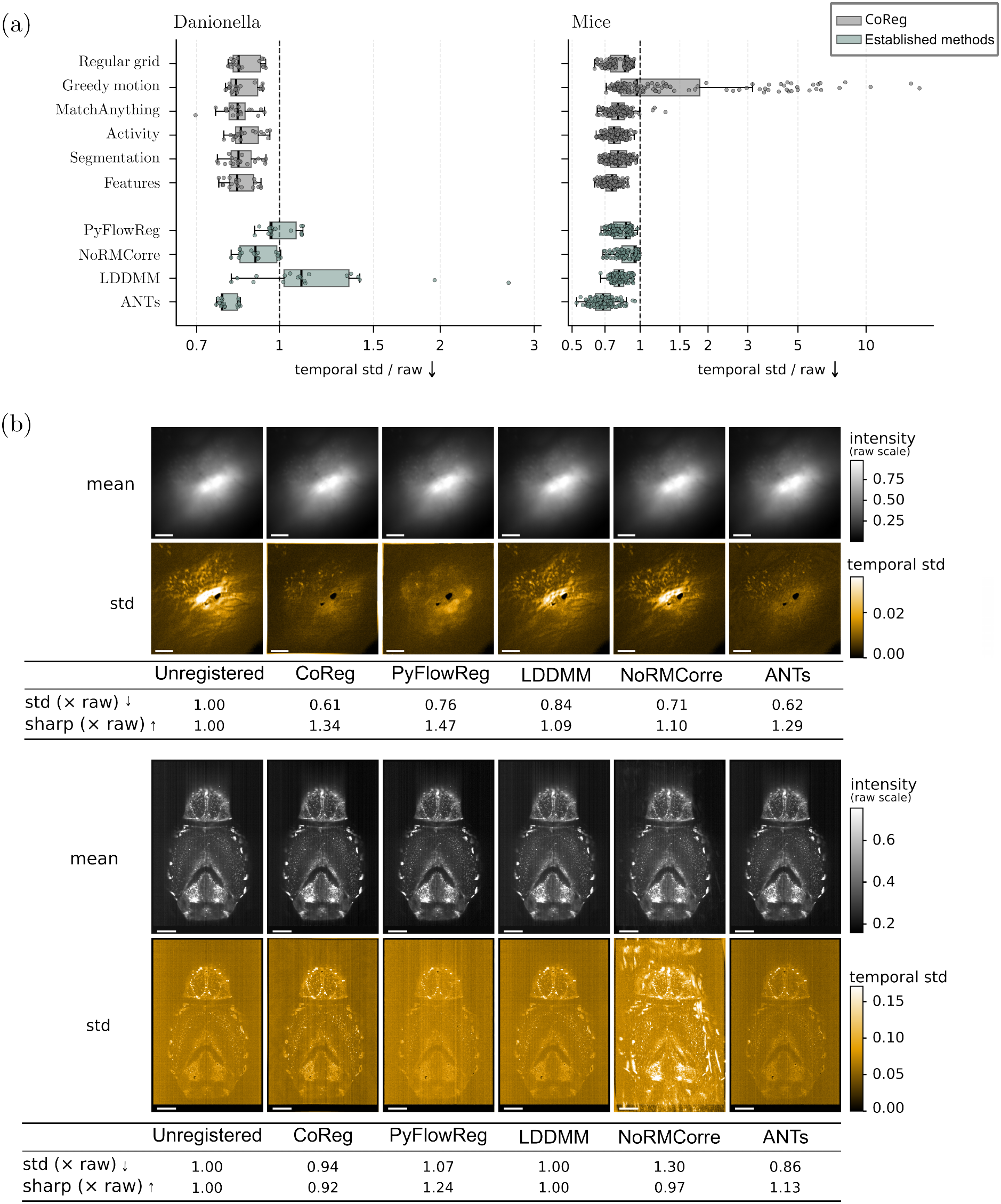
Motion compensation across time. (a) Temporal standard deviation (std) of each registered video divided by the std of the raw video, for the Danionella and mice datasets. The lower, the better the motion compensation. Dashed line visualizes raw video. Each dot represents one entire clip (< 700 frames) or one chunk of up to 700 frames, if the sequence length exceeds 1000 frames (mice *∼* 8000 frames). Methods are grouped into **CoReg**-based methods and established methods. Boxplots follow the same convention as Figure 2. (b) Two example clips (top: mice, Scale bar indicates 50 px [physical unit not available]; bottom: Danionella, Scale bar indicates 200 µm). For each method, the temporal mean image (upper row) and the temporal std image (lower row) are shown over all frames. In each example, both rows use one intensity scale across all methods. The tables provide the mean of the std image (“std”) and the sharpness of the mean image (“sharp”), both in relation to the raw image.

## Discussion

In this study, we treat motion correction in calcium imaging as a point tracking problem and introduce **CoReg**. Our results show that point tracking is a suitable framework for motion correction across different species and imaging setups. A key feature of our approach is the use of temporal information. Instead of aligning each frame independently to a static reference, **CoReg** tracks points across consecutive frames, which allows us to follow how motion develops over time. This addresses a key limitation of existing frame-by-frame methods, which estimate each deformation independently and cannot account for how motion evolves over time. As described in the Methods, **CoReg** uses surrounding frames to estimate point positions, producing trajectories that remain consistent across the full recording. This is particularly valuable in calcium imaging, where fluorescence changes driven by neural activity can be misinterpreted as motion when frames are processed individually. Our sequence length analysis supports this. **CoReg**’s advantage is most pronounced in longer recordings with gradually evolving motion. When motion is confined to brief, isolated events, the benefit of multi-frame tracking is expected to be smaller.

In calcium imaging, motion artifacts are caused by various factors. They can, for example, develop gradually over several frames or appear as isolated events. Using temporal information can therefore make the long-term correction more consistent across the full recording. **CoReg** relies on CoTracker3 for point tracking, which was trained on natural videos rather than biological microscopy data. The fact that it generalizes well to noisy, low-contrast calcium recordings suggests that the structural features used for tracking (local intensity patterns and temporal consistency) are shared across imaging domains. This is practically useful, as it means **CoReg** does not require domain-specific training data and can be applied directly without retraining. We also investigated whether fine-tuning CoTracker3 on calcium imaging data could further improve tracking performance using two adapter strategies. Both reduced the training loss, but the validation loss plateaued early, and quantitative metrics showed only marginal differences from the pretrained baseline. Fine-tuning, therefore, did not yield a meaningful improvement in registration quality (Supplementary Figure 7).

Simple translations are relatively easy to correct. However, many motion artifacts involve non-linear deformations, where different parts of the image undergo distinct local displacements. Correcting both types of motion is important for the method to be useful across different experimental conditions. We also find that performance is robust to the choice of tracked points, which makes the tool easier to use for researchers without a background in image processing. **CoReg** also offers several complementary advantages, such as using temporal information across frames, not requiring a fixed reference image, and allowing the tracked points to be inspected and corrected by the user. These properties make the correction process more interpretable and consistent across long recordings, even when frame-wise metrics do not clearly separate the methods.

Our results also highlight a divergence between frame-wise metrics and downstream analysis quality. PyFlowReg, for example, achieved high scores on metrics such as SSIM and ZNCC but performed poorly in preserving cell size and signal amplitude relative to the original video (see Fig. 2b, Fig. 4c, Fig. 4d). This suggests that standard registration metrics do not fully capture whether a method is suitable for downstream calcium analysis^43^. Registration-induced morphological distortion has been noted in other fluorescence imaging contexts^44^. Evaluating motion correction methods should therefore include qualitative checks and downstream measures such as cell area preservation and Δ*F/F*_0_ fidelity, not only frame-wise similarity scores. Future work could exploit the known temporal dynamics of GCaMP indicators as an additional quality criterion. Image changes faster than the indicator’s rise and decay times are unlikely to reflect true neural activity. They could therefore serve as a signal of residual motion or noise.

Better motion correction can improve signal extraction and the reproducibility of results. It may also reduce the number of recordings excluded due to strong motion artifacts. **CoReg** offers a temporally aware alternative to frame-by-frame methods. It is also more targeted than general-purpose registration frameworks that were not developed with calcium imaging in mind. One limitation is the dependence on reliable point tracking, which can be affected by low image quality, fast motion, or strong intensity changes. Motion that displaces tissue out of the imaged plane cannot be recovered by any planar registration method, including **CoReg**. Future work could integrate CoTracker3’s per-point confidence scores to flag frames or regions where tracking has likely failed, similar to bad-frame detection in Suite2p^9^. It could also include integration into live acquisition pipelines and extension to volumetric registration. By including both our method and the established methods in one tool, **CoReg** also aims to support more consistent and reproducible motion correction practices in the field. Therefore, **CoReg** provides a practical and transparent starting point for standardizing motion correction in calcium imaging research.

## Acknowledgements

This work was funded in part by the Deutsche Forschungsgemeinschaft (DFG, German Research Foundation) – Project numbers 554586006; 562675554; 531605337

## Author contributions statement

L.N. designed algorithms and experiments, implemented algorithms, analyzed and interpreted the data, and wrote the manuscript. M.P. improved algorithms and tested established methods. J.C.D., D.M., A.A., P.S., N.D., P.R., S.G. provided data, tested the CoReg platform, and interpreted the results. A.M.K. supervised the project, secured funding, and interpreted results. All authors reviewed the manuscript.

## Data availability

The data analyzed in this study are not publicly available but can be obtained from the corresponding author upon reasonable request. The code used for analysis is publicly available at https://github.com/ankilab/CoReg.

## Additional information

The authors declare no competing interests.

